# scATAcat: Cell-type annotation for scATAC-seq data

**DOI:** 10.1101/2024.01.24.577073

**Authors:** Aybuge Altay, Martin Vingron

## Abstract

Cells whose accessibility landscape has been profiled with scATAC-seq cannot readily be annotated to a particular cell type. In fact, annotating cell-types in scATAC-seq data is a challenging task since, unlike in scRNA-seq data, we lack knowledge of “marker regions” which could be used for cell-type annotation. Current annotation methods typically translate accessibility to expression space and rely on gene expression patterns. We propose a novel approach, scATAcat, that leverages characterized bulk ATAC-seq data as prototypes to annotate scATAC-seq data. To mitigate the inherent sparsity of single-cell data, we aggregate cells that belong to the same cluster and create pseudobulk. To demonstrate the feasibility of our approach we collected a number of datasets with respective annotations to quantify the results and evaluate performance for scATAcat. scATAcat is available as a python package at https://github.com/aybugealtay/scATAcat.

## INTRODUCTION

Chromatin structure can control the accessibility of potential gene regulatory elements in a dynamic and cell-type-specific manner and therefore plays a critical role in gene regulation (1). It has been shown that accessibility measurements offer valuable additional information to gene expression and have been demonstrated to be more cell-type specific than expression data (2). This is due to much of the cell-type-specific information within the genome being located in enhancer regions, which are captured by accessibility assays. Today, even single-cell technologies are applied to accessibility measurements. One of the most commonly used techniques is scATAC-seq (3). Further advances in single-cell genomics have facilitated the profiling of thousands of cells simultaneously, even at a multi-modal level (4). While these protocols lead to an enormous growth in the volume of data generated, the large datasets also allow deeper insights into complex biological systems.

Numerous types of cells can be found in an organism. Traditionally, these cell types have been defined based on phenotypic characteristics (5). With the advent of single cell RNA sequencing (scRNA-seq) it has become a common practice to cluster cells based on their transcriptome profile, in the expectation that these clusters correspond to cell types (6). The results, e.g., of the Human Cell Atlas (7), rely on this assumption and cell-types are frequently assigned to individual cells based on the determined transcriptome.

In this work we deal with the problem of assigning cell-types to the cells for which a single cell ATAC sequencing (scATAC-seq) experiment has been performed. This is an important problem if we want to exploit the detailed cell-type information that is assumed to be contained in the accessibility of the chromatin. Since most data and most cell-type annotation is available for scRNA-seq data, many existing approaches capitalize on the RNA level by predicting a proxy to gene expression from the accessibility landscape. This proxy is frequently referred to as “gene activity score” (8, 9). Given this gene activity score, annotation methods for scRNA-seq data can be carried over. These fall roughly into marker-based and reference-based methods. Marker-based annotation usually is a manual process which requires expert knowledge and testing of multiple markers. Recent work (10, 11) has tried to better support this process. Transferring labels from a known cell-type to the query cells by way of reference-based annotation can be achieved either by the statistical similarity metrics (12–16) or by machine learning models (17–21). (22) provides a benchmarking study comparing these techniques. A recent tool, Cellcano (23) is developed specifically for cell type annotation in scATAC-seq data and demonstrated superior performance over the existing approaches.

An alternative and potentially more accurate way to annotate cell types in scATAC-seq data is to perform within-modality annotation, that is to say, annotating the cell types using annotated-ATAC-seq data. Ideally, this will rely on an annotated scATAC-seq reference. One of the recent efforts to tackle this problem is EpiAnno (24) which leverages existing annotated scATAC-seq data and employs a Bayesian neural network framework for supervised cell-type annotation. However, this method is not computationally scalable (23). Often times the scATAC-seq reference itself is a product of annotation via expression markers on RNA levels. As a surrogate to annotated scATAC-seq references, an alternative approach is to use characterized-bulk ATAC-seq data as a reference. This approach has been suggested by (25).

Here we put forward *scATAcat - scATAC-seq cluster annotation tool* for annotation of cell-types in scATAC-seq data based on characterized bulk ATAC-seq data. scATAcat provides results comparable to or better than many approaches that rely on gene activity score. Rather than using the genes and their predicted activity as the features for assignment, we focus on the regulatory elements in the chromatin. We explore the use of FAC-sorted scATAC-seq data as a demonstration of the methodology. We further apply our annotation method and compare it to other approaches using PBMC and bone marrow sc-multiome data. For comparison we study two approaches, namely, marker-based annotation and reference-based label-transfer. We will discuss the challenges and biases in cell-type annotation in scATAC-seq data.

## MATERIALS AND METHODS

### scATAcat

#### Method outline

Given a set of cells for which a scATAC-seq experiment has been performed, scATAcat seeks to annotate cells with their corresponding cell-type. To be more precise, we first cluster the scATAC-seq data yielding what we call “pseudobulk” clusters. This serves the purpose of aggregating the sparse counts from several cells and to obtain better accessibility profiles at the pseudobulk level. All the cells in a pseudobulk cluster will inherit the assignment of the pseudobulk cluster. Furthermore, for annotation we require prototypes of the possible cell-types. Typically, for each cell-type there may be several replicate bulk samples available. Such prototypes can, e.g., come from existing characterized bulk ATAC-seq data. scATAcat takes as input these bulk accessibility profiles of distinct cell-types to which the computed pseudobulk clusters can be matched. scATAcat co-embeds the prototype accessibility profiles with the pseudobulk clusters in a principal component analysis (PCA) space and exploits the distance in this space to assign the cell-type labels.

#### Preprocessing of reference bulk ATAC-seq data

To allow for integration of different data sets we rely on candidate cis-regulatory elements (cCREs) provided by ENCyclopedia Of DNA Elements (ENCODE) project (26) as a feature space for all the datasets in this study. We calculate the cCRE coverage of the bulk ATAC-seq datasets. Next, we identify the differentially accessible cCREs in pairwise manner. This approach is adopted based on our hypothesis that these particular cCREs hold the most discriminative information and are cell-type specific. We use the DiffBind R package (v3.0) (27) with DESeq2 (28) as the underlying method after applying sequencing depth normalization provided by the package. We identify significantly differential accessible regions by filtering for FDR *<*= 0.05.

This analysis results in variable number of differential regions depending on the similarly between the compared cell types. To ensure equal contribution of various comparisons and mitigate the potential bias, we gather the same number of features, by default 2000 regions, from each pairwise comparison to derive a final feature space. When choosing this number we took into consideration the lower bound for the differential regions, as well as the total number of regions for the final feature set. Note that this number may be adjusted based on the specific cell-types under consideration. These features, hereafter named *differential cCREs*, are used as the final feature space for the rest of the analysis. We apply library size normalization and *log2* transformation to the original data and finally subset the matrix to *differential cCREs*.

#### Preprocessing of scATAC-seq data

Due to the sparsity of scATAC-seq data, some preproccessing is needed. For scATAC-seq data, we calculate the coverage of cCREs by counting the number of fragments within each cCRE region for each single cell. This yields a *cell-by-cCRE matrix*. Features which occur in less than *k* (default *k* = 3) cells get eliminated. Additionally, we get rid of the Y chromosome to avoid gender biases. On the level of cells, we filter out cells with fewer than 1000 and more than 80000 non-zero features, as well as doublet cells detected by AMULET (29).

Analysis suites for scATAC data (8, 9) process scATAC-seq data using TF-IDF. More specifically, we apply TF-logIDF normalization (30). This results in re-weighted features (cCREs) by assigning greater weight to more important features. Then we subset the data to the *differential cCREs* as defined by the reference bulk ATAC-seq data. We then reduce the dimension via PCA and continue with the standard scanpy clustering pipeline (31). We determine the nearest neighbors with *neighbors* function (n pcs = 50, n neighbors = 30)) and compute a UMAP (32) embedding. UMAP provides a low dimensional, non-linear embedding in which one can visualize a clustering. Next, we apply Leiden clustering (33) with *leiden* function.

The cells in one cluster form a *pseudobulk*. To represent this pseudobulk we compute its accessibility profile by adding the read coverage for each feature across the cluster member cells. Just like for the bulk data, we apply library size normalization and *log2* transformation to the pseudobulk matrix and subset the matrix to *differential cCREs*.

#### Co-embedding prototypes with pseudobulks

To make datasets comparable we apply a z-score transformation. First consider the *bulk-by-cCRE matrix* of prototypes. We substitute each matrix entry by its z-score with respect to its column, i.e. mean and standard deviation are computed over the cells in the column. Call this matrix *X*. For purposes of integrating the data, the z-score transformation on the *pseudobulk-by-cCRE matrix* is performed with the very same column-means and column-standard deviations of the first matrix. Call this transformed pseudobulk-by-cCRE matrix *Y*. Both matrices have as many columns as there were differential features. The number of rows, *n_X_*, of the transformed bulk-by-cCRE matrix is the number of annotated prototype bulk samples given. These may describe a number *k* of different cell-types. For one cell-type there may be several (replicate) bulk samples. The number of rows of *Y*, *n_Y_*, is just the number of pseudobulk clusters.

After this transformation step, we proceed to co-embed prototype bulk samples and pseudobulk clusters into one space using PCA. To this end, we obtain the eigenvectors of the matrix *X*. Depending on the number of dimensions one wants to keep, concatenate this number of eigenvectors into a matrix *W*. The data can then be represented in a lower dimensional space by transforming the original matrix as follows:

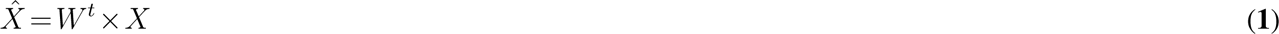

where *W^t^* corresponds to the transpose of the matrix *W*. The pseudobulk samples get projected onto the same PCA space as follows:

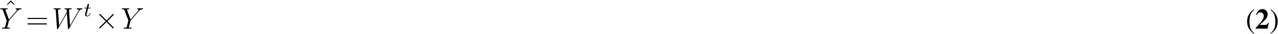

As a result, we have projection with both pseudobulk and prototype bulk samples embedded into the same space.

#### Annotating pseudobulk clusters

Co-embedding 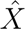 and 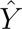, and visualizing the projection in 3D PCA space facilitates a simplified interpretation of cell-type relationships. However, determining the annotations solely based on a visualisation may be tedious. In particular, the first three PCs used in the projection might not suffice to fully capture the inherent structure of the high dimensional data. Following common practice, we therefore keep a larger number of dimension (typically 30 if there are enough samples available) to compute Euclidean distances in this high-dimensional embedding space.

To obtain a matching between pseudobulk clusters and prototype samples, we first compute centroids for each of the prototype cell-types. Next, we compute the Euclidean distances (in many dimensions) between the pseudobulk clusters and the *k* centroids, yielding a *n_Y_ × k* matrix *D*. Finally, we annotate each pseudobulk cluster by its closest centroid of a cell-type.

Going beyond the mere assignment, inspection of the matrix *D* can provide valuable information about the data. This will already show when decisions are ambiguous. Additionally, we also compute a matrix of size (*k* +*n_Y_*) *×* (*k* +*n_Y_*) with all the distances among pseudobulks and prototypes. Hierarchical clustering on this matrix helps to understand the grouping among pseudobulks and prototypes, as well as to detect possible outliers. scATAcat is implemented in Python and made compatible with scanpy (31) and anndata (34) libraries for an easy integration.

### Marker-based annotation

Marker-based annotation is one of the standard ways to annotate cell-types in scRNA-seq data. This method includes exploiting the expression of so-called marker genes, which are the genes exclusively expressed in a known cell-type, as a predictor of the associated cell-type. Application of this method for scATAC-seq data is enabled by the use of a metric called a “gene activity score” as a stand-in for the expression of genes. The gene activity score typically measures the level of accessibility around the gene body and nearby enhancer regions with the assumption that expression can be inferred from accessibility. Therefore, this value is also referred as *predicted* expression. In this study we use the gene score calculations as defined by Signac (8) which considers the number of fragments within the gene body and 2 kb upstream region of each gene to determine for each gene’s activity. Here we refer to the resulting matrix as *gene-score matrix*. Note that the scATAC-seq data is processed as outlined by Signac with default parameters to obtain this *gene-score matrix*. Once the gene activity scores are calculated, one can look at the predicted expression levels of the marker genes to determine the cell type of a cluster. In this work, CellMarker 2.0 (35) is used as the marker gene resource, if not specified otherwise,

### Reference-based label-transfer

Another commonly used way to annotate cells in single-cell data includes *transferring* the cell labels from existing annotated reference atlases. Here we use one of the widely adapted tools, Seurat (v3) (13, 36) for this purpose. Briefly, Seurat (v3) employs canonical correlation analysis (CCA) to determine so-called “anchors” between the query and reference datasets. Anchors represent the cells with the highest similarity and therefore serves as a correspondence between reference and the query. The assumption is that the anchor cells define matching cell states and create a shared space between the query and reference. Each anchor pair is then scored based on the correspondence in connecting cells’ shared nearest neighbor (SNN) graphs (37). Anchors serve as bridges to *transfer* information, such as cell-type label, from reference to query cells.

It is important to note that in order for reference and query to align and for bridges to be defined, (i) there needs to be a shared feature space between the reference and query and (ii) both the data need to be processed similarly. To satisfy the first requirement, we use *gene-score matrix* of scATAC-seq data described above, effectively defining our features as genes. For the second requirement, we apply *logTransformation* for both the datasets. We next determine the variable features using Seurat’s *vst* method and select top 3000 variable features. Subsequently, we determine anchors using *FindTransferAnchors* method with “cca” as the reduction method. Note that, in our experience, the latest Seurat4 integration with reduction method “spca” typically results in limited and insufficient number of anchors. Consequently, we chose to use “cca” reduction method to attain more informed annotations. Lastly, we use *TransferData* function to transfer cell-type information from reference to query data. We apply this approach for cell-type annotation, which we referred to as *label-transfer*.

### Performance assessment

In the assessment of annotation methods, we adapted the performance metrics introduced in (38). Given an experiment with a set of cells, each cell have a ground truth annotation, whereby each cell is assigned a label (e.g., T-cell). While each method provides an annotation, the annotations may not cover all cells, and the labels may not encompass all the labels that are used by the ground truth annotation. Let the set of annotation labels be *L* = 1,2*,…,K*. Further, let *C* be the set of cells *c_i_*. We only use cells that have been annotated by the ground truth and by all methods which we compare. Let *m_i_*denote method *i*, where we adopt the convention that *m*_1_ is the ground truth. *m_i_* is a mapping from cells to labels, i.e. it is of the form *m_i_*(*c_j_*) = *l* with *l ∈L*. This would mean that mapping *m_i_* labels cell *j* as type *l*. Note that we only use those labels that are predicted by all methods, i.e. we reduce the label set to 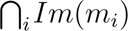. Accordingly, we also reduce the cells in the ground truth and only keep those cells for which the label is a member of this reduced label set. The first metrics we use, *accuracy* (*Acc*), quantifies the proportion of cells with accurately assigned cell-type annotations across cells and calculated as

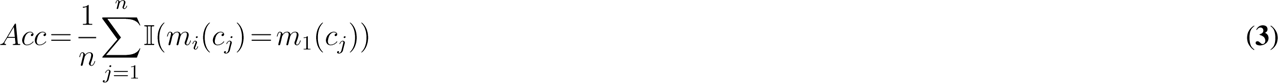

where *n* is the number of labeled cells, *c_j_*is the *j^th^*cell, *m_i_* is the mapping from cell to predicted cell-type and *m*_1_ is the mapping from cell to true (ground-truth) cell-type.

*Balanced accuracy* (*BAcc*) measures the average accuracy across cell-types and calculated as:

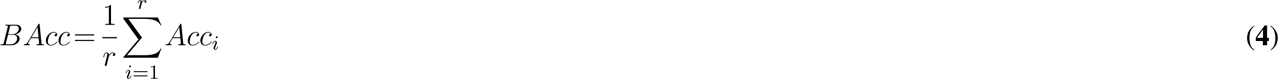

In this equation *r* denotes the number of cell-types in reduced cell-type subset 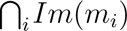. *Acc_i_*refers to the accuracy of the *i^th^* cell-type.

The last metric we use is called *cluster accuracy* (*CAcc*). The clusters are defined by scATAcat pipeline using the Leiden clustering as explained above (see Section *Preprocessing of scATAC-seq data*). For each annotation method, as well as the ground-truth, every cluster is annotated as the most abundant cell-type of its constituent cells. We only consider the clusters with more than 10 cells. Let *p_i_*denote method *i*, where we adopt the convention that *p*_1_ is the ground truth cluster labels. *p_i_* is a mapping from clusters to labels, i.e. it is of the form *p_i_*(*s_j_*) = *l* with *l ∈L*. This would mean that mapping *p_i_* labels cluster *j* as type *l*. The cluster accuracy is calculated as follows:

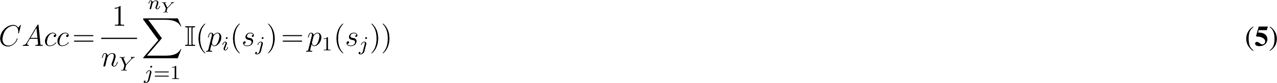

where *n_Y_*refers to the number of clusters as defined by the Leiden clustering, *s_j_* is the *j^th^* cluster, *p_i_* is the mapping from clusters to predicted cell-type and *p*_1_ is the mapping from clusters to true (ground-truth) cell-type.

### Datasets

#### scATAC-seq datasets

##### FACS human hematopoiesis (bone marrow) scATAC-seq

The bone marrow scATAC-seq data is acquired from (39). The dataset includes scATAC-seq data of 2,210 cells human hematopoietic progenitor cells which are obtained from bone marrow and isolated via fluorescence-activated cell sorting (FACS). FACS enables sorting of single cells into a well plate using cell-type specific cell-surface markers and thereby provides a true cell-type annotation for each cell in the data.

Owing to this unique property, we use this data both as single-cell data as well as bulk prototype data. When using as a bulk prototype, we combine the single cells of the same cell-type by summing the features across cells and form *bulk-like* data. As the data is by default annotated, here we refer to this dataset as annotated bulk prototype dataset. When using as a single-cell data, as the name suggests, we use the data obtained from cells individually and do not leverage the FACS annotations.

##### Human PBMC sc-multiomics

Human PBMC sc-multiomic data has been obtained from 10X website^1^. The dataset includes paired scRNA-seq and scATAC-seq profiles of 10,661 cells, obtained from a peripheral blood mononuclear cells (PBMCs) of a healthy donor. Inference of the cell-type annotations of this dataset is introduced in the Methods section.

##### Human bone marrow sc-multiomics

This dataset has been provided as part of the NeurIPS challenge (40). The dataset serves as a valuable resource for benchmarking studies and claimed to be the largest and most realistic multimodal data available (41). The datasets are also carefully annotated by experts considering various marker genes and across scRNA-seq and scATAC-seq data. We focused on one sample from this study, namely, *s1d1* for the sake of simplicity. The data consist of 19,039 bone marrow cells obtained from a healthy donor. In this study, we decoupled the paired measurements and exclusively used the scATAC-seq data. For this dataset, we use the annotations provided by the challenge organizers as the ground-truth, therefore, do not perform cell-type annotation for the scRNA-seq part.

#### Reference datasets

##### Human PBMC CITE-seq

Human PBMC CITE-seq (36) comprises the transcriptomic measurements of 211,000 PBMCs along with 228 cell-surface proteins. The reference includes two levels of cell-type annotations with increasing granularity, namely, cell-type.l1 and celltype.l2. We used the coarse annotation, celltype.l1, which includes the main blood cell types, namely; B, CD4 T, CD8 T, natural killer, monocytes, dentritic, other T and other cells.

##### Human BMNC CITE-seq

The BMNC (bone marrow mononuclear cell) dataset (13) comprises single-cell transciptomics measurements of 33,454 bone marrow cells along with 25 cell-surface proteins. This reference as well, includes two levels of cell-type annotations with increasing granularity, namely, cell-type.l1 and celltype.l2. The used fine-grained annotation, celltype.l2, which includes progenitor cells and hematopoietic stem cells, along with the PBMCs.

##### Human hematopoietic differentiation bulk ATAC-seq

The human hematopoietic differentiation bulk ATAC-seq dataset (2) includes bulk ATAC-seq profiles of 13 human primary blood cell-types extending over diverse hematopoiesis layers. Samples are obtained from peripheral blood and bone marrow. In our study, we exploit distinct subsets of this data to match potential cell-types in the given query dataset. As the FACS-profiled scATAC-seq bone marrow data inherently specifies the cell-types, we match the bulk prototypes to those cell-types. When integrating the bulk samples with the sc-multiome bone marrow data we use all 13 cell-types. Lastly, when integrating with PBMC data, we consider only the terminal cell states (unipotent cells), which constitutes the largest portion of a typical PBMC samples.

Notably, this dataset lacks the dendritic cell population. To account for this, we merge the the dataset with the plasmacytoid dendritic cells data from (42). All the data is preprocessed according to ENCODE ATAC-seq analysis pipeline (43).

Summary of the datasets and in which combination they are used for different methods is summarized in Supplementary figure 4.

## RESULTS

### Testing scATAcat on FACS bone marrow scATAC-seq data

To assess the performances of the cell-type annotation methods, we first used FACS-profiled scATAC-seq data. This part of the study serves as a feasibility study as both the query and the references essentially consist of the same dataset. The data is inherently annotated by cell-types due to the cell-surface markers of sorted cells. We leveraged these annotations to assess the efficacy of our scATAcat method. scATAcat requires bulk prototype data to provide cell-type annotation for query scATAC-seq data. We created *bulk-like* prototypes by aggregating the same cell-types from the sorted scATAC-seq. DiffBind requires minimum two replicate per cell-type to provide reasonable statistics regarding differential regions. Therefore, for each cell-type we created two bulk-like prototypes by randomly splitting the cells per cell-type into two. We next identified differentially accessible regions using DiffBind.

The cell-type commitments in early hematopoiesis follow hierarchical transitions (44). Based on data from (2, 42), we constructed such hematopoiesis tree shown in Figure 1A. We considered this differentiation trajectory when determining differentially accessible regions. More precisely, we defined the cell-type pairs for the detection of differential regions by considering *parent* - *child* relationship in the hematopoiesis tree. This enables adapting the features to the given system.

**Figure 1.**
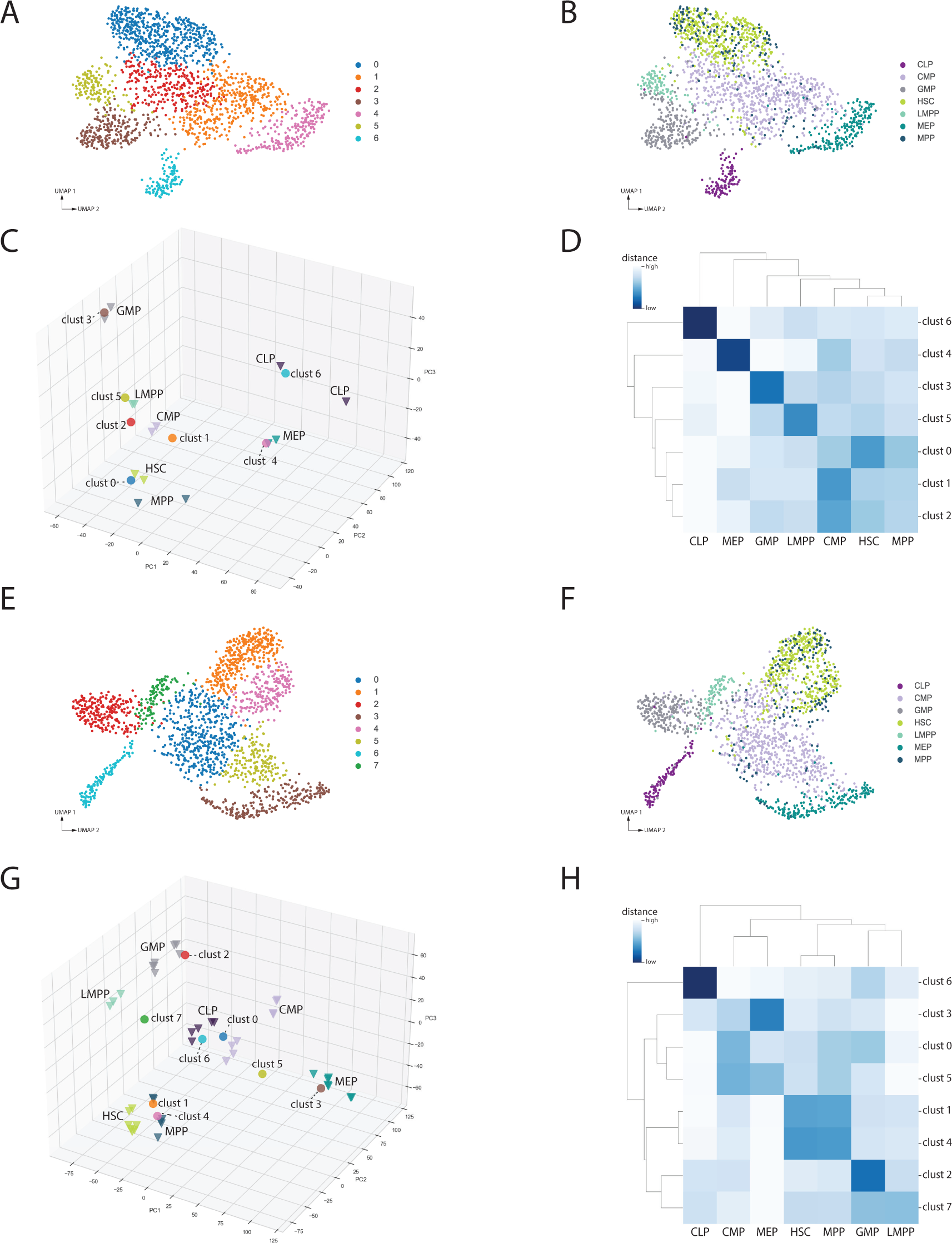
**(A-D)** Testing scATAcat on FACS bone marrow scATAC-seq data as part of the feasibility study. **(A)** UMAP embedding colored by the clustering of FACS scATAC-seq cells, and **(B)** by true (ground-truth) cell identities derived from FACS labels of common lymphoid progenitor (CLP), common myeloid progenitor (CMP), granulocyte-monocyte progenitor (GMP), hematopoietic stem cell (HSC), lymphoid-primed multipotent progenitor (LMPP), megakaryocytic-erythroid progenitor (MEP) and multipotent progenitor (MPP). **(C)** 3D PCA projection of the feasibility study. Prototypes, depicted as triangles, are created by aggregating single cells from (B) based on cell-type, with two prototypes generated for each cell-type. Circles represent pseudobulks, the aggregated forms of clusters from (A). **(D)** Heatmap of the high-dimensional Euclidean distances between prototypes and pseudobulks. **(E-H)** Testing scATAcat on FACS bone marrow scATAC-seq data with bulk ATAC-seq prorotypes. **(E)** UMAP embedding colored by the clustering of FACS scATAC-seq cells, and **(F)** by true (ground-truth) cell identities derived from FACS labels. **(G)** 3D PCA projection of the pseudobulks together with prototype bulk samples. Triangles represent prototypes while circles represent pseudobulks. **(H)** Heatmap of the high-dimensional Euclidean distances between prototypes and pseudobulks.

Figure 1A shows the clustering of the single cells into 7 clusters (Leiden resolution 0.65). From this clustering we compute 7 pseudobulk samples. The clustering is not in full agreement with annotation, as can be seen in Figure 1B. Hematopoietic stem cells (HSCs) and multipotent progenitors (MPPs) do not show a clear separation and multipotent common myeloid progenitors (CMPs) are clustered into two subgroups. Based on the annotation, we computed bulk-like prototypes. Next, we integrate the pseudobulks from the clustering with the prototypes in one PCA space. Figure 1C represents the 3D PCA projection obtained through scATAcat. The visualization clearly illustrates a distinct separation among prototypes. The pseudobulk clusters closely align with these prototypes.

While the 3D projection provides an intuition about the relationships between prototypes and pseudobulks, the Euclidean distances from high dimensions provide more accurate proximity information. Therefore, we determined the Euclidean distances between the prototypes and pseudobulk samples in high dimensional PCA space consisting of 30 principal components (PCs). A heatmap representation of these distances is depicted in Figure 1D. For the purpose of assignment, each pseudobulk gets assigned to its closest prototype. Accordingly, Cluster 0 corresponds to HSC; Cluster 1 and 2 to CMP; Cluster 3 to granulocyte-monocyte progenitor (GMP); Cluster 4 to megakaryocytic-erythroid progenitor (MEP); Cluster 5 to lymphoid-primed multipotent progenitor (LMPP); and Cluster 6 to common lymphoid progenitor (CLP).

When one now carries over this annotation to the single cells that make up the pseudobulk one can count the percentage of correctly annotated cells. We use three metrics to assess the accuracy including *accuracy*, *balanced accuracy*, and *cluster accuracy*. Accuracy represents the percentage of correctly assigned cells while balanced accuracy averages these calculations across cell-types. In the current example, accuracy is 81% and *balanced accuracy* reaches to 78%. The last metric, cluster accuracy, measures the accuracy across clusters. In this feasibility study, all the clusters are correctly annotated with 100% cluster accuracy.

The matrix of Euclidean distances between pseudobulks and prototypes already shows that some decisions are almost a tie. An alternative visualization is provided by clustering all the distances among pseudobulks and prototypes. Such a heatmap with the associated single-linkage dendrogram is shown in Figure S1B. In this figure, e.g., the group of cluster 1, 2, and the CMP cells form a cluster in the dendrogram.

After demonstrating the effectiveness of our method in the feasibility study, we applied scATAcat again to the same FACS scATAC-seq data. However, this time, we integrated external prototypes from bulk ATAC-seq data (2). These prototypes align with the cell-types present in the scATAC-seq data. We define differentially accessible regions using the prototypes and use these features to represent the data. Therefore, the data representations differ despite using the same scATAC-seq data.

Figure 1E shows the clustering (Leiden resolution 0.6) of cells in a UMAP embedding, revealing 8 clusters. We form pseudobulks by aggregating the cells in each cluster. Figure 1F depicts the real identities of the cells. Clusters show remarkable consistency with annotations. Nevertheless, HSCs and MPPs do not exhibit a clear separation and CMPs are again clustered into two subgroups. Our next step was to co-embed pseudobulks with prototypes. Figure 1G depicts the 3D PCA projection produced by scATAcat. The figure contains colored triangles for the prototype replicates and circles for the pseudobulks. Mostly there is a clear association between the correct prototypes and pseudobulks recognizable, although there is larger variability among the CMP and GMP prototypes. HSCs and MPPs are less clearly distinguishable but mix with each other, as can also be seen in the UMAP.

Subsequently, we compute the Euclidean distances between the prototypes and pseudobulks in high dimension to compute more informative cell-type annotation. Figure 1H shows the heatmap of these distances. Using this distance matrix we annotate each pseudobulk by the closest prototype cell-type. In this way we can correctly associate clusters 0 and 5 to CMP, clusters 1 and 4 to HSC, cluster 2 to GMP, cluster 3 to MEP, cluster 6 to CLP, and cluster 7 to LMPP. This shows that using the high-dimensional euclidean distances scATAcat again provides accurate annotations for each cluster.

We next applied marker-based annotation to the same FACS scATAC-seq data and aimed at annotating cell-types by the *predicted* expression profiles of the marker genes in cells. Marker-based annotation requires collection of cell-type specific marker genes which in itself may be a nontrivial task. We obtain marker genes from a curated data table from (45) (Table S3). As these genes are by definition cell-type specific, we expect also their predicted expression to be specific for the same cell-type. Figure 2 shows the UMAP embedding of the cells with the color code depicting ground-truth cell identities for HSCs and CMPs (A and E, respectively). However, as shown in Figure 2 B-D for HSCs (respectively Figure 2 F-H for CMPs) the marker genes do not show the distinctive predicted expression profiles. In fact, in both cases marker genes are expressed in a dispersed set of cells over the UMAP. Some marker genes show ubiquitous predicted expression (2C) while others show unspecific characteristics. We conclude that transcriptome based marker genes cannot be directly carried over to predicted gene expression space to annotate cell-types in scATAC-seq data.

**Figure 2.**
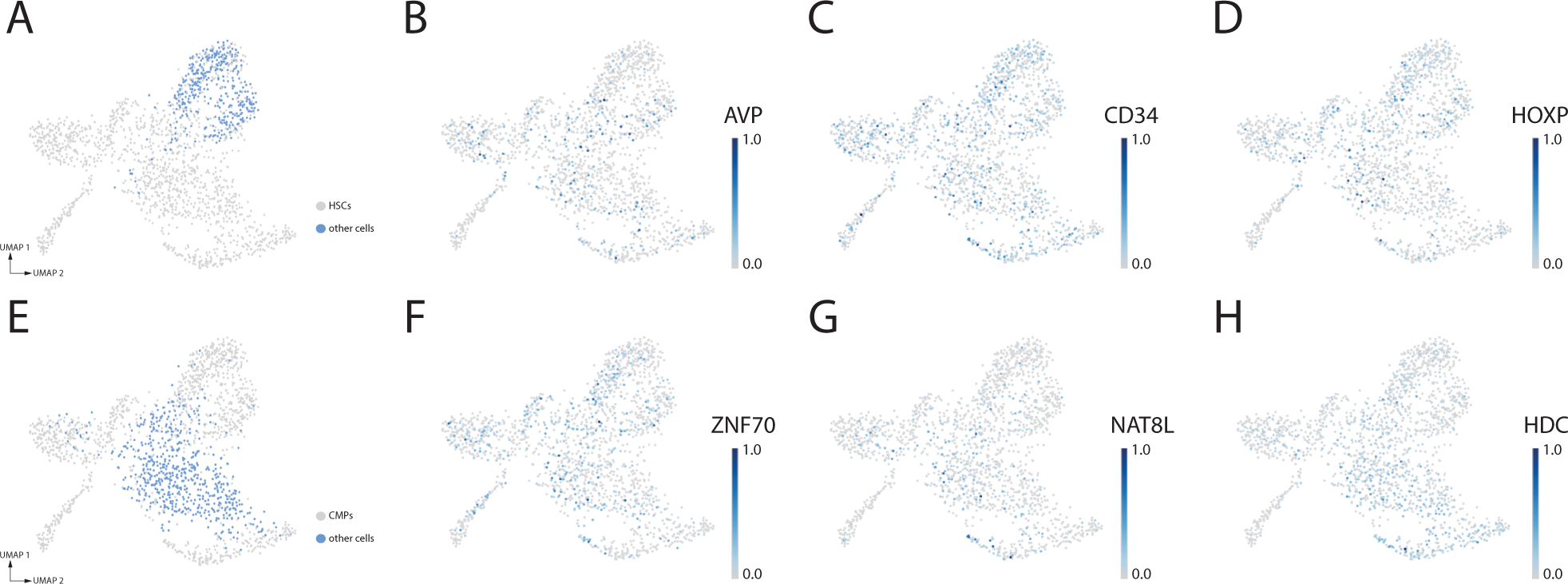
Marker-based annotation of hematopoietic stem cells (HSCs) and common myeloid progenitors (CMPs). (A) depicts UMAP embedding of FACS-characterized bone marrow scATAC-seq data. The blue colored cells represents ground-truth HSCs. The figures (B-D) show the predicted expression levels of HSC marker genes, AVP, CD34 and HOXP, with darker shades indicating higher expression. (E) depicts UMAP embedding of FACS bone marrow scATAC-seq data. The blue colored cells represents ground-truth common myeloid progenitors (CMPs). The figures (F-H) depict the predicted expression levels of CMP marker genes, ZNF70, NAT8L, HDC, with darker shades indicating higher expression.

Lastly, we employed Seurat’s label-transfer approach on the same FACS-characterized bone marrow scATAC-seq data for cell-type annotation. Label-transfer essentially uses Canonical Correlation Analysis to transfer cell-type labels from a well-annotated reference dataset to a query dataset. In order to use this approach we rely on the BMNC CITE-seq data (see Datasets) as a reference and predicted gene expression data of FACS bone marrow scATAC-seq data as the query. Since the CITE-seq data includes measurements of cell-surface proteins, the cells in the reference dataset have reliable and independent annotations. Figure 3C shows the UMAP embedding of the scRNA-seq part of reference CITE-seq data which serves as the source for transferred cell-type labels. Figure 3A depicts the UMAP representation of the query scATAC-seq data and the color code depicts the real cell-type identities. Some of the cell-types form a distinct cluster, e.g. CLPs, GMPs. However, some cell-types display inconsistency between the clustering and their real cell-type identities. Notably, HSCs and MPPs cannot be distinguished. Additionally, HSC and MPPs, as well as MEPs are split into two clusters. Finally, Figure 3B shows the annotated FACS bone marrow scATAC-seq data where the color code denotes cell-type annotations obtained with the label-transfer approach. These annotations do not show uniformity within clusters and annotations do not overlap with the original cell-type identities.

**Figure 3.**
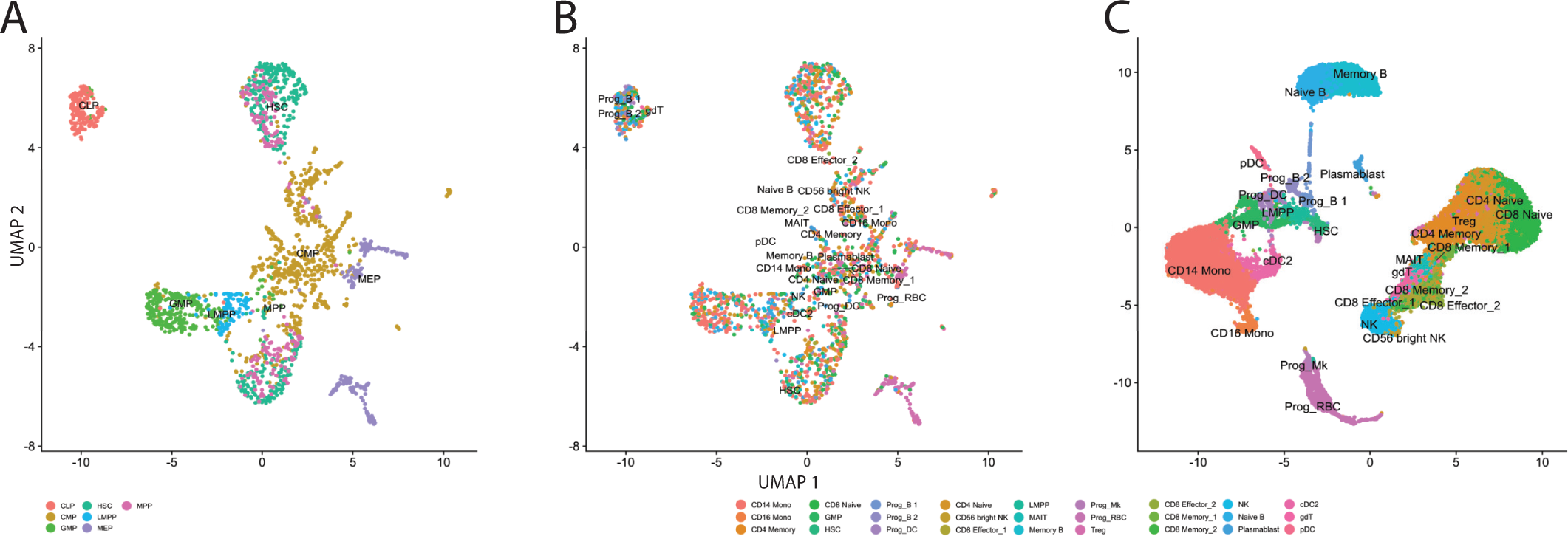
Cell-type annotation via label-transfer. **(A)** UMAP representation of predicted gene expression matrix colored by ground-truth cell-types. **(B)** shows the same UMAP embedding however, this time the color-code indicates the predicted cell-type identity by the label-transfer approach as shown by the legend. **(C)** scRNA-seq UMAP embedding of the reference bone marrow data, used as a reference when applying label-transfer approach, colored by cell-type identities.

In order to quantitatively assess this, we compare the annotations obtained from the label-transfer approach to the real cell-type identities obtained from FACS labels. The annotations in reference BMNC CITE-seq data cover a broader spectrum of bone marrow cells than those of query FACS labels. Additionally, the annotations of reference and query datasets disagree on parts of the nomenclature, even though they may correspond to similar cell types. To increase the concordance between annotations, we modified the annotations to a coarser annotation scheme whenever possible, for example, we used ‘monocyte’ instead of ‘CD14 Monocyte’. Similarly, nomenclature is aligned between the annotations, for example, we used ‘CLP’ instead of ‘B cell progenitor (Prog B)’. For the annotations that did not allow such simplification (e.g., ‘ILC,’ ‘gdT,’ and ‘cDC2’), we retained their original cell identities. Original annotations along with their corresponding simplified annotations are provided in Supplementary Table 1. The common cell-type annotations between CITE-seq data and FACS labels are MEP, GMP, HSC, CLP and LMPP. Therefore we focus on these common annotations. We assessed the accuracy using three metrics; *accuracy*, *balanced accuracy*, and *cluster accuracy*. The label-transfer approach correctly annotates 16% of the cells while on the same cell-type subset scATAcat demonstrates a significantly higher accuracy of 92%. This trend is consistent in both balanced accuracy, where scATAcat achieves 92% compared to the label-transfer approach’s 20%, and cluster accuracy, with scATAcat at 86% and the label-transfer approach at 14%.

Since the granularity of Leiden clustering results depends on the resolution parameter, we tested the impact of this parameter on the performance of scATAcat and its resulting comparison to label-transfer. We applied scATAcat with Leiden clustering resolutions varying from 0.1 to 2.5. This was done on the same FACS bone marrow scATAC-seq data and the prototypes introduced above for scATAcat. For the label-transfer approach the settings and the data are the same as above. We compared the resulting annotations with ground-truth cell identities obtained from FACS labels. Accuracy metrics, including *accuracy* (*Acc*), *balanced accuracy* (*BAcc*), and *cluster accuracy* (*CAcc*) are presented in Supplementary Table 3. Both accuracy and balanced accuracy show fairly robust values across varying clustering resolutions. Cluster accuracy is inherently affected by the number and the homogeneity of clusters, as reflected in our results.

### Performance of scATAcat in annotating PBMC data

Single cell transcriptome technologies have been frequently demonstrated and used on PBMCs and thus provided deep understanding of blood cell-types. For purposes of this study it is particularly helpful that a sc-multiome PBMC dataset is available combining nuclear scATAC-seq with cytoplasmic scRNA-seq of the same cells (see Datasets).

Since the PBMC sc-multiome data has not been externally annotated, we are leveraging the scRNA-seq part of the data to create a surrogate ground truth. Relying on the assumption that cell-type annotation in scRNA-seq data is more straightforward (46) than for scATAC-seq, we can evaluate our scATAcat annotation with respect to this ground truth.

To obtain a ground truth cell type annotation, scRNA-seq data gets processed using Seurat’s standard pipeline (36). Subsequently, cell-type annotation is performed via the label-transfer approach using PBMC CITE-seq data as the reference and applying *SCTransform* for both the reference CITE-seq and query scRNA-seq data. Anchors are determined using *spca* as the reduction method and cell-type identities are extracted via the *MapQuery*. The cell-type annotations of the scRNA-seq data is shown in Figure S2A.

Having a ground-truth annotation of the cells available, we proceeded to annotate the scATAC-seq part of the multiome data using scATAcat, via the marker-genes, and by label-transfer. To apply scATAcat, scATAC-seq data is processed as introduced earlier. For the prototypes we again use the FACS-characterized hematopoietic bulk ATAC-seq data (see Datasets), albeit now focusing only on those terminal cell states (CD4 T cells, CD8 T cells, B cells, NK cells, monocytes, dendritic cells) that are also part of PBMCs (47). The differentially accessible regions between the bulk prototypes are identified and used as the reference frame for clustering. A UMAP representation of the clustering (Leiden resolution 1.0) of the cells is shown in Figure 4A. We formed 15 pseudobulks out of these clusters and applied scATAcat to integrate prototypes and pseudobulks.

**Figure 4.**
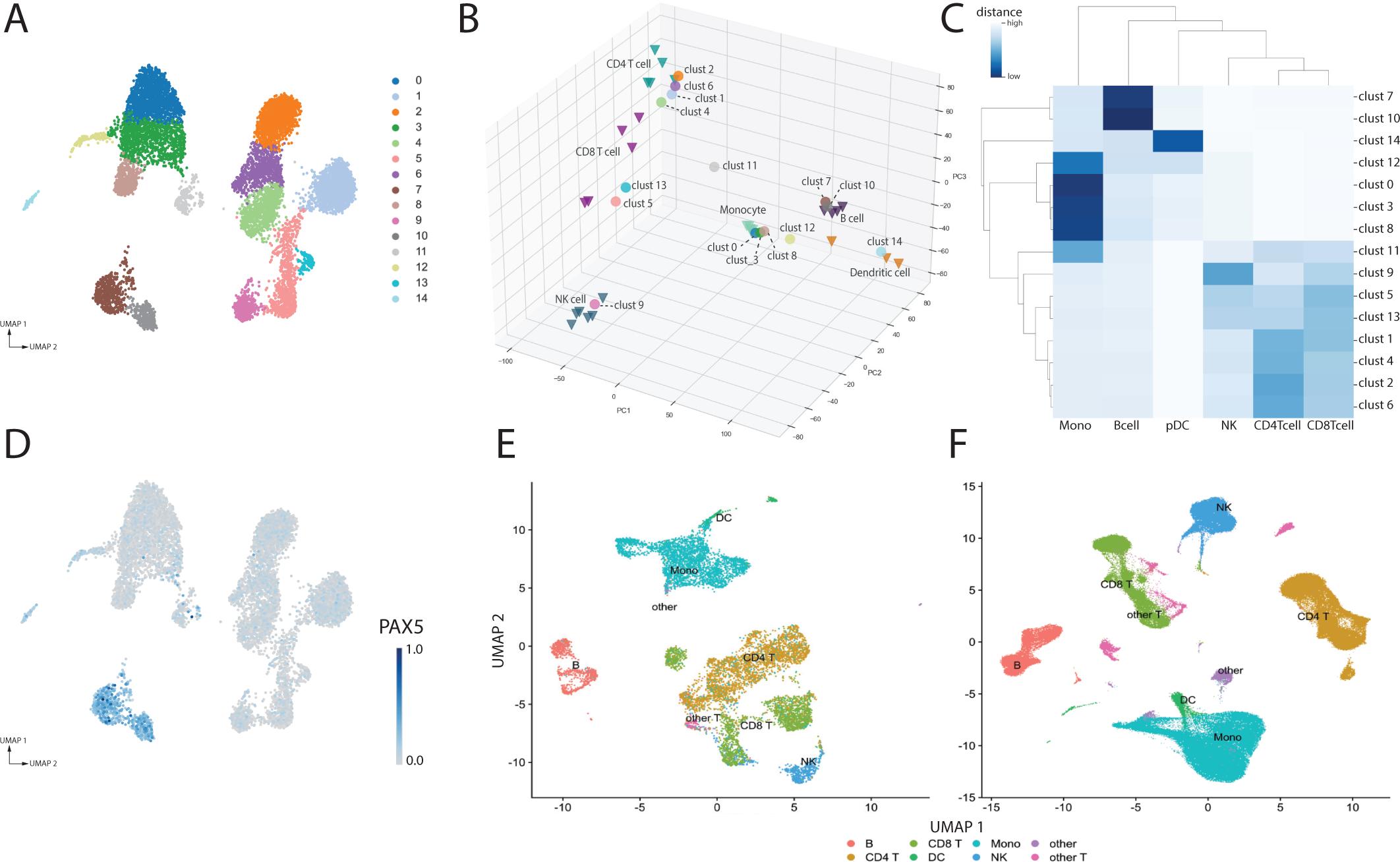
Performance of scATAcat in annotating PBMC data. **(A)** UMAP embedding colored by the clustering of PBMC scATAC-seq cells. **(B)** 3D PCA projection of the PBMC scATAC-seq pseudobulks together with prototypes. Triangles represent prototypes while circles represent pseudobulks. **(C)** Heatmap of the high-dimensional (50 PCs) Euclidean distances between the pseudobulks and prototypes shown in (B). Monocyte (Mono), B cell (Bcell), plasmacytoid dendritic cell (pDC), natural killer cell (NK), CD4 T cell (CD4Tcell), CD8 T cell (CD8Tcell). **(D)** UMAP embedding PBMC scATAC-seq cells colored by predicted marker gene expression of B cell marker PAX5. **(E-F)** Cell-type annotation via label-transfer. **(E)** UMAP representation of predicted gene expression matrix colored by predicted cell-type labels based on label-transfer. **(F)** scRNA-seq UMAP embedding of the reference PBMC data, used when applying label-transfer, colored by cell-type identities. Other cell types (Other), other T cell subtypes (other T).

Figure 4B depicts the 3D PCA projection obtained by scATAcat. The bulk prototypes roughly cluster into three groups, akin to the UMAP plot. Pseudobulks closely align and co-embed with bulk samples except for cluster 11. The replicates of CD8 T cell and dendritic cells are more scattered in 3D space indicating a higher variability among them. Figure 4C shows the heatmap of the high-dimensional Euclidean distances between bulk prototypes and pseudobulks. The heatmap depicts a clear clustering structure for most of the clusters resulting in clear annotations. However, the high similarity between CD4 and CD8 T cells (also visible in Figure S2B) makes it harder to annotate the clusters surrounding them. Nevertheless, we annotated each pseudobulk by the closest bulk sample, hence, Clusters 0, 3, 8, 11, and 12 correspond to monocyte; cluster 1, 2, 4 and 6 to CD4 T cell; cluster 5 and 13 to CD8 T cell; cluster 7 and 10 to B cell; cluster 9 to natural killer (NK) cell; cluster 14 to plasmacytoid dendritic cells (pDC).

Although cluster 11 is annotated as a monocyte, it shows similarity with other prototypes, too. This observation could be attributed to a biological characteristic, such as the presence of highly similar cell types in the scATAC-seq data. In such intricate cases, it is helpful to further investigate the data. An alternative visualization provided as part of scATAcat, depicting the clustering of pairwise Euclidean distances between pseudobulks and prototypes, can be helpful in interpreting such cases. Figure S2B shows the heatmap of these distances. Cluster 11 shows affinity to both the monocyte prototype together with other monocyte pseudobulks (clusters 0, 3, 8, 12) as well as to the CD4 T-cells with clusters 1, 2, 4 and 6. Possible explanations for this phenomenon include heterogeneity in cluster 11 or possible doublet cells. The annotation process will respect the closest distances in high dimensions and annotate cluster 11 as monocytes. Yet, the example shows that visual inspection of the high-dimensional distances can indicate potential biological or technical problems.

To put these results in context, we generated the predicted expression levels of PBMC scATAC-seq cells and carried out marker-based annotation. Predicted expression levels of a well-established B cell marker gene, PAX5 (48), is depicted in Figure 4D. Cluster 7 and 10 show highest expression of PAX5 suggesting these clusters as potential B cells. Notably, we also observe a low-level but ubiquitous expression of PAX5 across other cell types. Figure S2D-I illustrates further examples of the marker-based annotation which further demonstrates the non-quantitative property of this annotation approach.

As a final annotation strategy, we employed the Seurat’s label-transfer approach to the scATAC-seq part of the multiome data. We used the same procedure as in the label-transfer for the FACS bone marrow scATAC-seq data, although now based on PBMC CITE-seq data as reference. As a query we use the predicted expression values of PBMC scATAC-seq data. Figure 4E shows the predicted annotations of the query cells and Figure 4F represents the scRNA-seq UMAP embedding of the reference CITE-seq data. Annotations shows high concordance with clusters, although dendritic cells appear close to monocytes indicating a lack of separation. Similarly, CD4 T cells cluster closely with CD8 T cells. This pattern is further reflected in cell-type annotation - some of the cells forming a cluster with CD8 T cells are annotated as CD4 T cells or NK cells. Likewise, certain cells within the CD4 T cell cluster are annotated as monocytes.

Lastly, we compared the annotations obtained from scATAcat and label-transfer. We excluded the marker-based annotation from this comparison because choice of marker genes is subjective and making clear decisions about the annotation is difficult. As indicated in 4F, the PBMC CITE-seq reference includes the additional labels “other T cells” and “others”. Thus, we ignore these cells and consider only the cells annotated with a common cell-type that is shared with the bulk prototypes. Based on the ground truth derived from the RNA part of the multiome data, scATAcat correctly annotated 84% of the cells while label-transfer correctly annotated 91% of the cells. When these annotations are averaged across cell-types, i.e. balanced accuracy, scATAcat reached to 77% while label-transfer achieved 90%. This consistent pattern is also evident in cluster accuracy, where scATAcat performed at 85% and label-transfer achieved 92%. These results are robust across various clustering parameters as shown in Supplementary Table 4.

### Performance of scATAcat in annotating bone marrow sc-multiomoe data

Lastly, we applied the three annotation strategies to bone marrow sc-multiome data (see Section Datasets). Like in sc-multiome data, we focus on the scATAC-seq part of the data and simply ignore the scRNA-seq part. This dataset has been provided as part of a benchmarking competition and it comes with expert cell-type annotation. We first applied our scATAcat method for cell-type annotation. As prototype, we used both the progenitor and terminal cell types in the bulk data and determined differentially accessible regions between pairs of cell-types. For the progenitor cell-types, we incorporated the hematopoiesis trajectory by selecting the pairs of cell-types based on their parent-child relationships within the lineage tree shown in Figure S1A. For the leaf nodes of the tree, i.e. the terminal cell states, we conducted pairwise comparisons between the cell types that share a common progenitor. Up-to 2000 most differential regions per comparison get combined to obtain the final feature set. Then we preprocess the scATAC-seq data and perform clustering.

A UMAP of the clustered cells (Leiden resolution 1.0) is shown in Figure 5A. We form 12 pseudobulks out of these clusters and integrate them with prototypes using scATAcat. The result of this projection is depicted in Figure 5B. Most of the pseudobulk-clusters project closer to terminal cell-types than to progenitors. Besides, as expected by their high prevalence in bone marrow (49), many clusters project closer to erythrocytes. Most of the lymphoid cells (CD8 T cell, CD4 T cells and NK cells) show rather indistinctive positioning in the 3D projection. Figure 5C shows a heatmap of the high-dimensional Euclidean distances between prototypes and pseudobulks.

**Figure 5.**
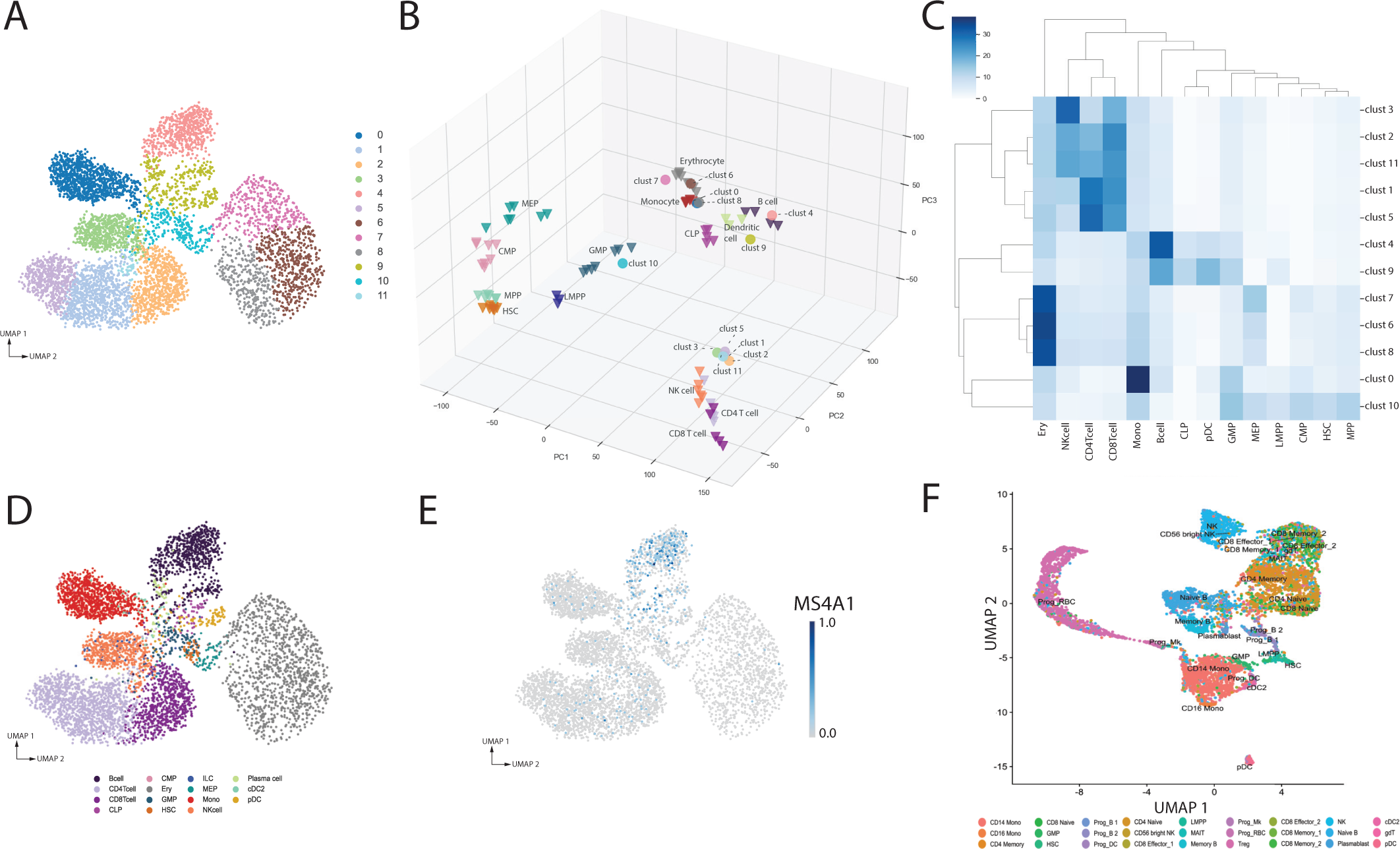
Performance of scATAcat in annotating bone marrow data. **(A)** UMAP embedding of scATAC-seq cells of bone marrow sc-multiome data colored by the clustering. **(B)** 3D PCA projection of the bone marrow scATAC-seq pseudobulks together with prototypes. Triangles represents bulk samples while circles reprresent pseudobulk-clusters. **(C)** Heatmap of the high-dimensional Euclidean distances between the pseudobulks and prototypes shown in (B). Monocyte (Mono), B cell (Bcell), plasmacytoid dendritic cell (pDC), natural killer cell (NK), CD4 T cell (CD4Tcell), CD8 T cell (CD8Tcell), erythrocyte (Ery), common lymphoid progenitor (CLP), common myeloid progenitor (CMP), granulocyte-monocyte progenitor (GMP), hematopoietic stem cell (HSC), lymphoid-primed multipotent progenitor (LMPP), megakaryocytic-erythroid progenitor (MEP) and multipotent progenitor (MPP). **(D)** UMAP embedding of scATAC-seq cells of bone marrow sc-multiome data colored by the ground-truth cell-type identities. Innate lymphoid cells (ILCs), Conventional dendritic cells 2 (cCD2). **(E)** Cell-type annotation via marker-genes. UMAP embedding of scATAC-seq cells colored by predicted marker gene expression of B cell marker MS4A1. **(F)** Cell type annotation via label-transfer. UMAP representation of predicted gene expression matrix colored by predicted cell-type labels.

This representation of the distances shows a similar profile to 3D projection. The progenitors form a separate cluster and the lymphoid cells, while not well separated in the 3D PCA, form a cluster in the heatmap. Pseudobulks 1, 2, 3, 5, and 11 are nearby the prototypes for CD4 T cells, CD8 T cells NK cells in the 3D PCA and are clearly clustered together with them in the heatmap. In fact, this is expected considering the lineage tree shown in Figure S1A. We again annotate each pseudobulk by considering its closest prototype cell-type in terms of high-dimensional Euclidean distance. Cluster 0 corresponds to monocyte, clusters 1 and 5 to CD4 T cell; clusters 2 and 11 to CD8 T cells; cluster 3 to NK cell; cluster 4 and 9 to B cell; clusters 6, 7 and 8 to Erythrocyte; and finally, cluster 10 to GMP.

Next, we generated the predicted expression levels of the scATAC-seq profiles and employed marker-based annotation. We use the same marker genes here as we used in annotating FACS-profiled bone marrow scATAC-seq data (Figure 2). While progenitor cell-type marker genes only provided a fuzzy picture of cell identity, terminal cell-types display better predictive performance for the sc-multiome bone marrow data studied now (Figure S3). For example, B-cell marker gene MS4A1 clearly highlights the B-cell cluster (4 and 9) in the UMAP. Nevertheless, marker-based annotation remains the most subjective approach.

As a last cell-type annotation method, we carried out label-transfer approach. We followed the same procedure as employed in label-transfer for the FACS bone marrow scATAC-seq data. The BMNC CITE-seq data is used as the reference and predicted expression levels of sc-multiome (scATAC-seq) data as the query. Figure 5F shows the predicted labels of the query cells in the UMAP embedding. Generally, clusters are predominantly annotated by one cell type, although the annotations get mixed near cluster boundaries. Also, subsets of CD4 and CD8 T cell co-embed without a clear separation.

Next, we compare the cell-type annotations obtained via the scATAcat and label-transfer approach to the provided ground-truth annotations. All three annotations include similar cell-types at varying granularity. To increase the overlap between different annotation schemes and provide more inclusive comparison we simplified the annotations to a coarser annotation scheme, if possible. Original cell-type annotations along with their corresponding simplified annotations are provided in Supplementary Table 2. In the subset of cell-types shared between scATAcat, label-transfer and ground truth annotations, scATAcat demonstrated an accuracy of 96%, outperforming label-transfer at 79%. scATAcat also exhibited superior balanced accuracy, achieving 95% compared to label-transfer’s 71%. Similarly, scATAcat surpassed label-transfer with 89% accuracy, while label-transfer achieved 78%. Overview of these accuracy metrics across datasets are presented in Figure S4B. We examined the potential impact of the clustering parameter choice on accuracy. As presented in Supplementary Table 5, both accuracy and balanced accuracy are robust across varying clustering parameters while, as expected, cluster accuracy shows slight variations.

## DISCUSSION

Studying the accessibility profiles of cells enables understanding of the regulatory landscape governing each cell type. The current single-cell methods like scATAC-seq hold the promise of unraveling the gene regulation at a more precise resolution. However, the resulting data presents a number of challenges which need to be tackled to fully capitalize on the valuable of information they offer.

One of the foremost challenges in scATAC-seq data analysis is to annotate cell-types. Typically, the methods developed for cell-type annotation in scRNA-seq data are borrowed to annotate the cells in scATAC-seq data. This is enabled by transforming the scATAC-seq data into scRNA-seq-like-data using the accessibility around the genes as a proxy for expression. In this study, we have demonstrated that this approach may not yield optimal results, and does not fully exploit the potential of scATAC-seq, which inherently offers more cell-type specificity. We argue that annotation within the same modality would improve the cell-type annotations. To facilitate this, we developed scATAcat which enables cell-type annotation of clusters in scATAC-seq data by leveraging prototype accessibility profiles typically obtained from bulk ATAC-seq data.

Another challenge in scATAC-seq data is the sparsity. Limited information in individual cells makes it impractical to perform annotation at this level. As a remedy, we propose the pseudobulk. Given that the clustering is performed sufficiently fine-grained to not merge different cell-types, pseudobulk reflects the accessibility of the cell-type appropriately. Consequently, all the cells in a pseudobulk share the same cell-type annotation.

Another key decision in the design of our method was to replace the ATAC-peaks by ENCODE cCREs. As ENCODE cCREs are derived through the integration of various experiments across cell types, the advantage here is the stability of the reference frame which is a prerequisite for data integration across different experiments.

In addition to cell-type annotation, our program scATAcat provides useful visualizations for the interpretation of the these annotations. In one, we provide the heatmap of the Euclidean distances which illustrates the similarity between all the clusters and prototypes. Additionally, we provide a bipartite heatmap showing the similarities of pseudobulks to each prototype cell-type. This heatmap serves as a quantitative indicator, providing a measure of confidence in the annotations. In this way, one can, e.g., identify a potential doublet clusters showing similarity to multiple cell-types, as well as a new cell-type with poor annotation confidence to all the prototypes.

In our study, we observed a superior performance of scATACcat over the label-transfer approach in two different bone marrow datasets. Given that the bone marrow is where hematopoiesis takes place to produce various blood cell types, the bone marrow cells follow a differentiation trajectory and are expected to show a dynamic and heterogeneous accessibility profile (39). It has been suggested that during lineage commitment, *chromatin accessibility precedes gene expression* (19), suggesting more pervasive and steady accessibility profiles than expression profiles. We think that the good performance of scATACcat, which is based on the annotation in accessibility space, can be explained by this phenomenon.

Additionally, the superior performance of scATAcat was more pronounced in FACS-characterised scATAC-seq data in comparison to sc-multiome (scATAC-seq) bone marrow data. One reason for this can be the higher throughput of the sc-multiome data enabled by the recent droplet-based technologies. Nevertheless, having an external annotation in FACS-characterised scATAC-seq data provides a solid base for comparisons.

Interestingly, we observed an opposite trend in PBMC data. Unlike bone marrow cells, PBMCs mostly consist of nave or resting cells in the absence of effector functions (47). This suggests a stable expression pattern which can be better exploited by the gene-centric annotation strategies. Another explanation to the poorer performance of scATAcat on PBMC data can be the choice of the ground-truth annotations. In the absence of an external annotation for the this dataset, we annotated the scRNA-seq part of the multiome data with the label-transfer approach and used these annotations as the ground-truth. Establishing the ground-truth by label-transfer using the same reference data may introduce a bias in favor of this method in comparison to scATAcat.

While our method has provided valuable insights, it also comes with limitations. First of all, the scATACcat requires prototype cell profiles data as input, consequently, as in any reference based tool, can only annotate the cell-types for provided prototypes. Secondly, we built on the assumption that the pseudobulk clusters are sufficiently pure. However, this assumption may not hold true especially in the case of complex samples including similar cell-types. In general, we recommend opting for a higher number of clusters to ensure more homogeneous clusters. That being said, we also provided evidence that our method is fairly robust to clustering resolutions (Supplementary Tables 3, 4 and 5).

A potential further extension of the method can be enabled by the use of single-cell-pseudobulk prototypes instead of bulk prototypes. This approach would leverage the increasingly available and more homogenous scATAC-seq atlases. Additionally, conceptually our method is not restricted to scATAC-seq data. We are currently working on applying it to single-cell epigenetic data like single-cell DNA methylation or single-cell ChIP-seq.

## DATA AVAILABILITY

scATAcat is available as a python package at https://github.com/aybugealtay/scATAcat. The scripts used to generate the figures from this manuscript along with the figures/tables from the Supplementary materials and their respective outputs can be accessed at https://github.com/aybugealtay/scATAcat paper.

## ACKNOWLEDGMENTS

The authors would like to thank Zhicheng Ji for his valuable inputs regarding feature selection step. The authors also gratefully acknowledge the IT group of the Max Planck Institute for Molecular Genetics for their support and flawless maintenance of the computational resources.

---

**Supplementary figure 1.**
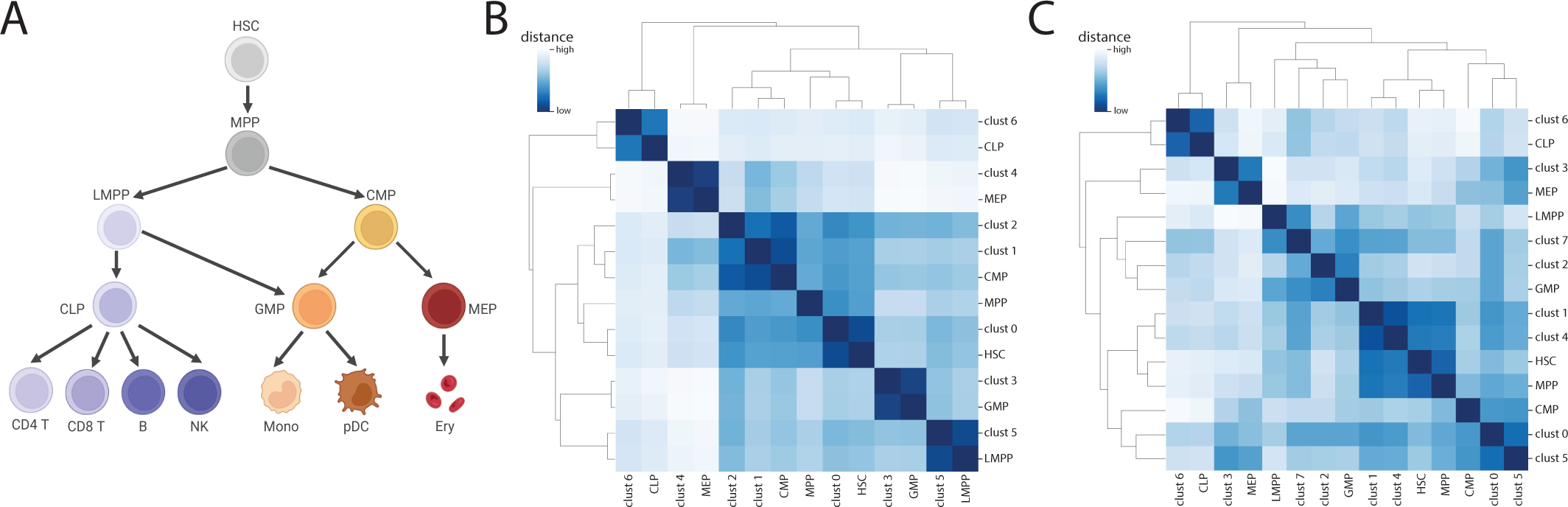
**(A)** Haematopoiesis tree depicting the cell-type differentiation. The tree is constructed for the cell-types included in this study. Monocyte (Mono), B cell (Bcell), plasmacytoid dendritic cell (pDC), natural killer cell (NK), CD4 T cell (CD4Tcell), CD8 T cell (CD8Tcell), erythrocyte (Ery), common lymphoid progenitor (CLP), common myeloid progenitor (CMP), granulocyte-monocyte progenitor (GMP), hematopoietic stem cell (HSC), lymphoid-primed multipotent progenitor (LMPP), megakaryocytic-erythroid progenitor (MEP) and multipotent progenitor (MPP). **(B)** A heatmap depicting the Euclidean distances among pseudobulks and prototypes in the feasibility obtained via scATAcat using FACS bone marrow scATAC-seq data. **(C)** A heatmap depicting the Euclidean distances among pseudobulks and prototypes obtained via scATAcat using FACS bone marrow scATAC-seq data and progenitor bulk prototypes.

**Supplementary figure 2.**
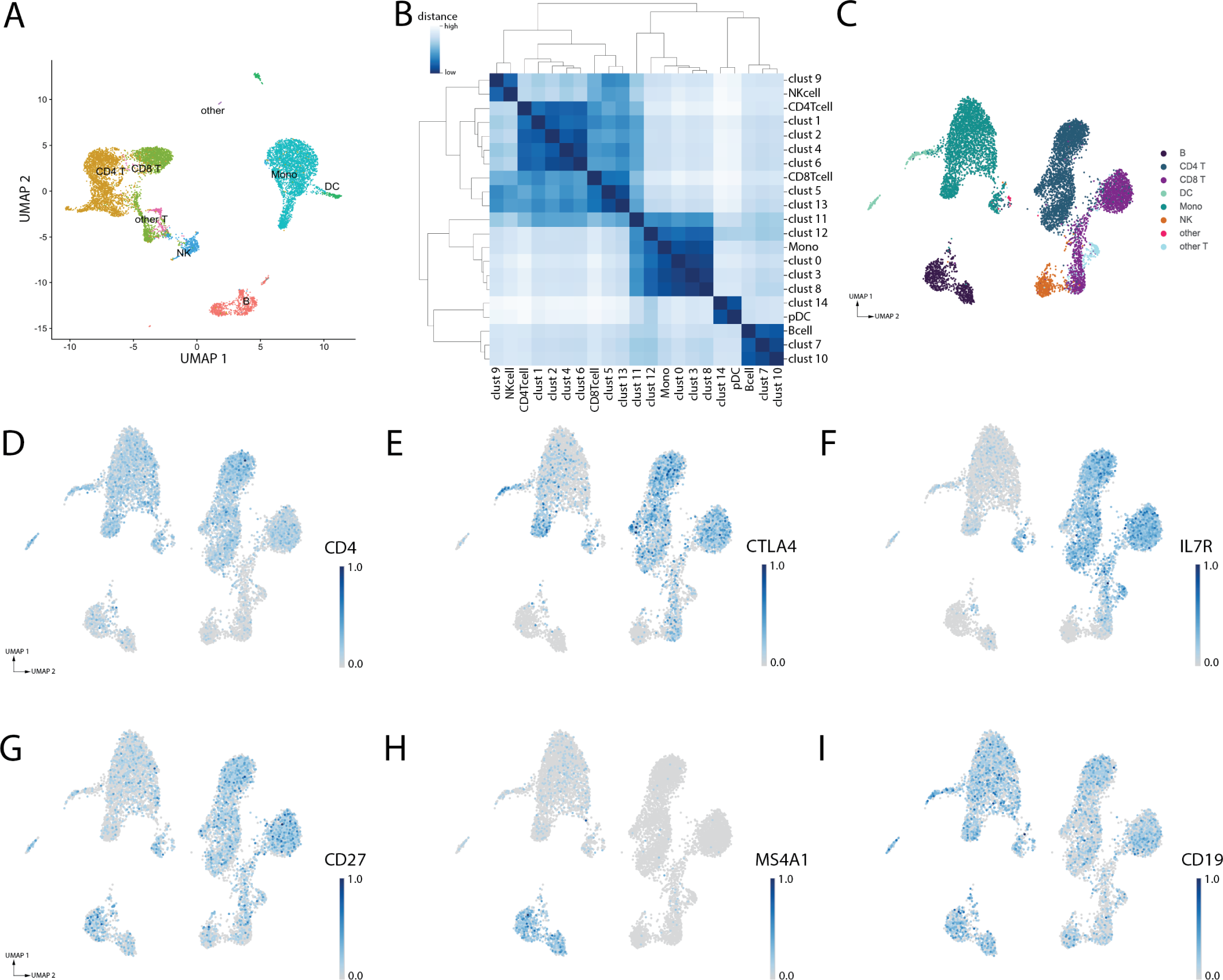
**(A)** UMAP visualization of scRNA-seq PBMC data (part of PBMC sc-multiome) depicting color-coded cell-type labels predicted using label-transfer. Monocyte (Mono), B cell (Bcell), dendritic cell (DC), natural killer cell (NK), CD4 T cell (CD4Tcell), CD8 T cell (CD8Tcell),other cell types (Other); other T cell subtypes (other T). **(B)** A heatmap depicting the Euclidean distances among pseudobulks and prototypes obtained via scATAcat using PBMC scATAC-seq data and bulk prototypes of PBMC cell-types. Plasmacytoid dendritic cell (pDC). **(C)** UMAP embedding of PBMC scATAC-seq data colored by the ground-truth cell-type identities. **(D-I)** Predicted expression levels of marker genes, CD4, CTLA4 and IL7R for CD4 T cells; CD27, MS4A1 and CD19 for B cells.

**Supplementary figure 3.**
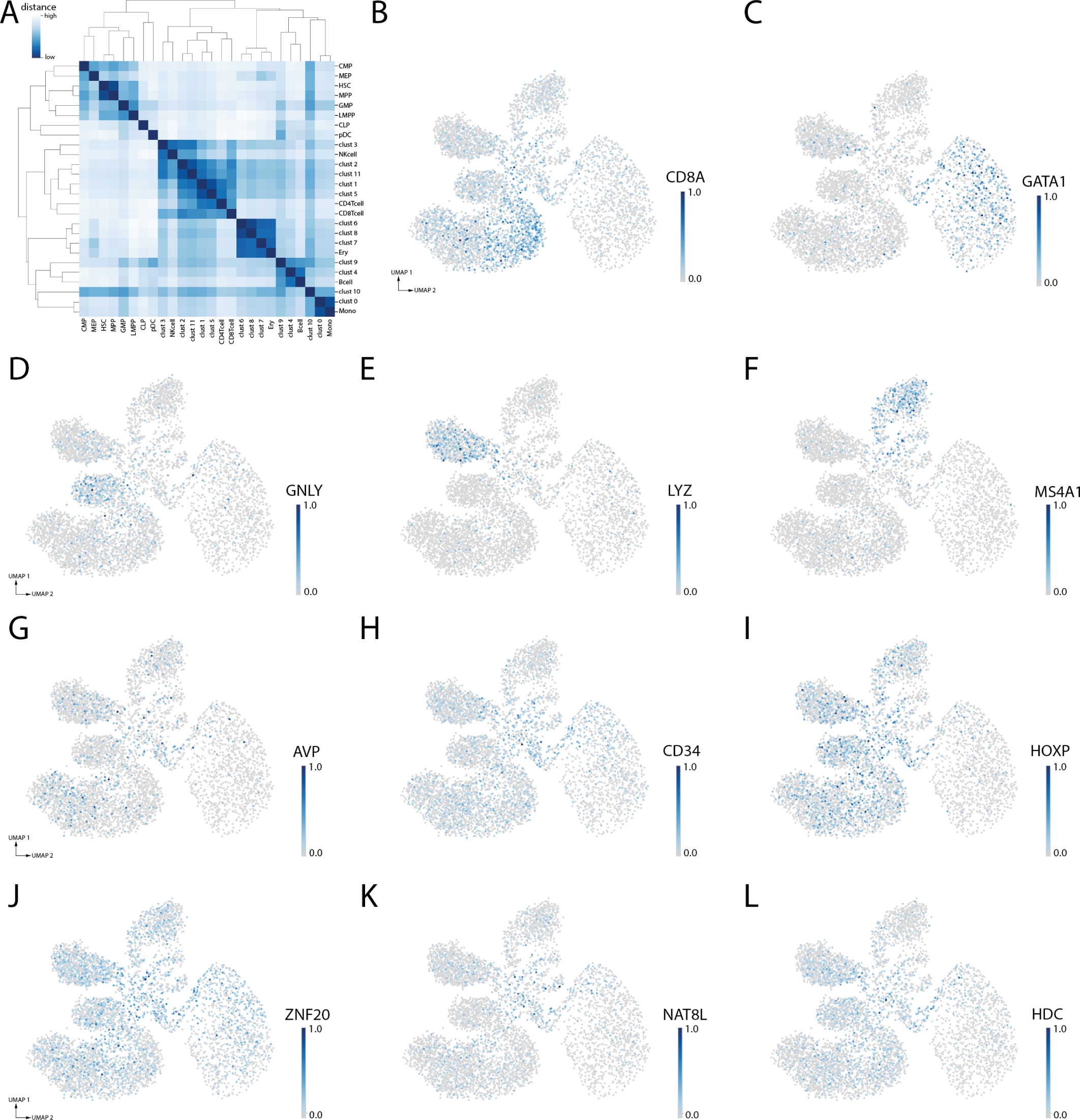
**(A)** heatmap depicting the Euclidean distances among pseudobulks and prototypes obtained via scATAcat using bone marrow scATAC-seq data (bone marrow sc-multiome) and bulk prototypes of progenitor and terminal cell-types (all cells in 1A). Monocyte (Mono), B cell (Bcell), plasmacytoid dendritic cell (pDC), natural killer cell (NK), CD4 T cell (CD4Tcell), CD8 T cell (CD8Tcell), erythrocyte (Ery), common lymphoid progenitor (CLP), common myeloid progenitor (CMP), granulocyte-monocyte progenitor (GMP), hematopoietic stem cell (HSC), lymphoid-primed multipotent progenitor (LMPP), megakaryocytic-erythroid progenitor (MEP) and multipotent progenitor (MPP). **(B-M)** Predicted expression levels of various marker genes, CD8A for CD8 T cells, GATA1 for erythrocytes, GLNY for natural killer cells, LYZ for monocytes, MS4A1 for B cells, AVP, CD34 and HOXP for hematopoietic stem cells, ZNF70, NAT8L and HDC for common myeloid progenitors. Darker colors indicate higher predicted expression.

**Supplementary figure 4.**
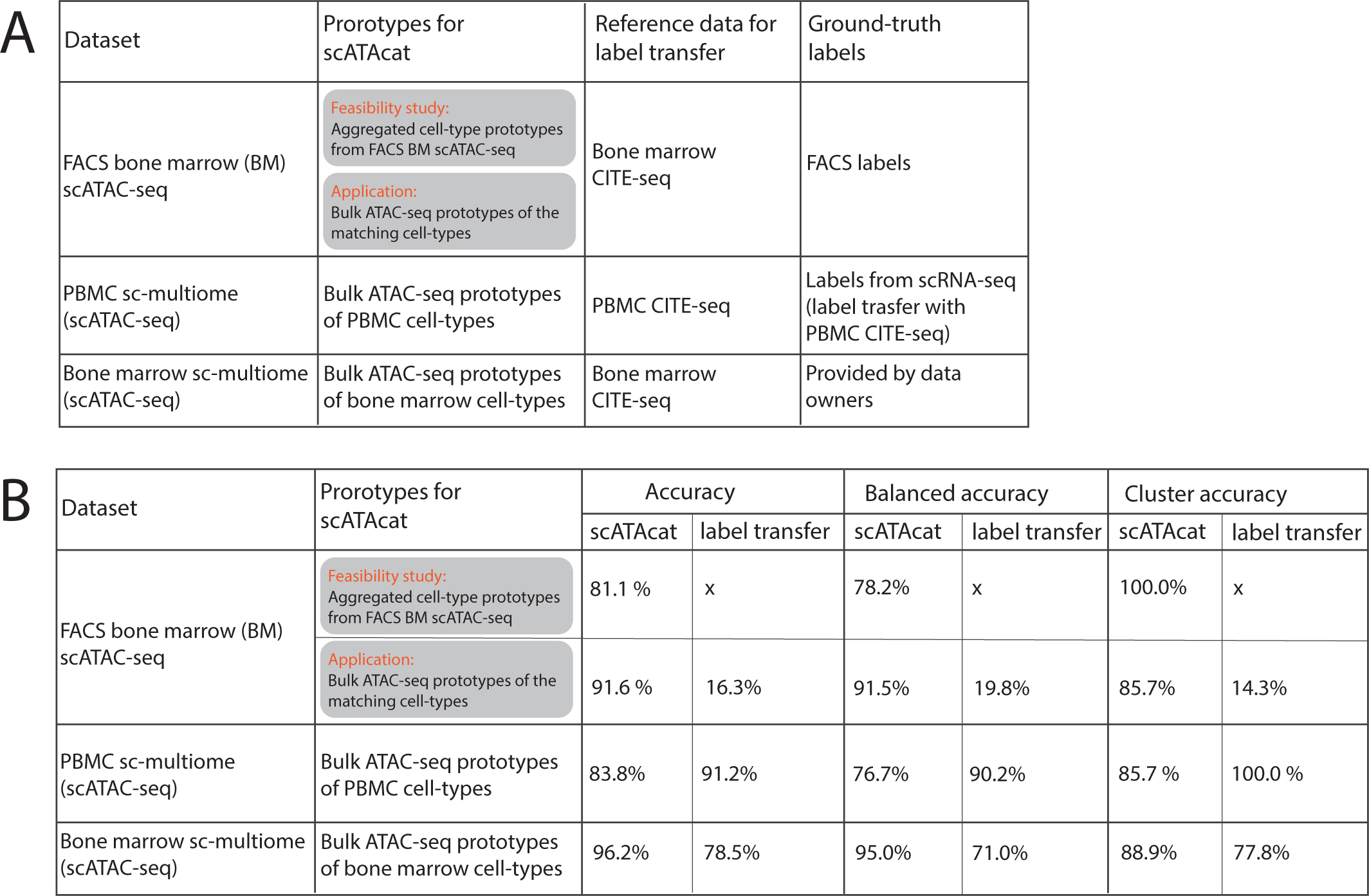
(A) Table summarising the datasets and methods used in this study. (B) Summary table presenting accuracy calculations for all datasets employed in this study, comparing across different methods.

## Footnotes

1 https://www.10xgenomics.com/resources/datasets/pbmc-from-a-healthy-donor-granulocytes-removed-through-cell-sorting-10-k-1-standard-1-0-0

